# Auxin and cytokinin antagonistically regulate Arabidopsis root distal stem cell fate through opposing pairs of HD-Zip II transcription factors

**DOI:** 10.64898/2026.10.23.760923

**Authors:** Giovanna Sessa, Marco Possenti, Ilaria Brozzi, Gianmarco Serrapica, Alberico Di Pinto, Altea Alfè, Massimiliano Sassi

## Abstract

The antagonistic action of auxin (IAA) and cytokinin (CK) regulates several aspects of plant development including stem cell specification and maintenance. In the *Arabidopsis thaliana* root, the fate of distal stem cells (DSCs) is mainly regulated by IAA, while the function of CK is still ill-defined. Here, by investigating the role of the HD-Zip II transcription factor family in distal tissues, we uncover CK as a proliferative input for DSCs, acting in opposition to IAA. We show that opposing pairs of HD-Zip II proteins regulate DSCs fate by integrating IAA and CK signals. The δ module, composed of HAT3 and ATHB4, is activated by CK and promotes DSCs division. Conversely, the γ module, constituted by HAT1 and ATHB2, acts downstream of IAA to repress CK biosynthesis and DSCs division. The mutual inhibition between δ and γ modules downstream of opposite hormone inputs sustains the precise regulation of DSCs homeostasis.

## INTRODUCTION

Indeterminate growth is one of the distinctive features of plants and constitutes the basis of their developmental plasticity^1,2^. The capacity of plants to generate new organs throughout their life cycle is granted by the stem cells contained within meristematic tissues and is governed by intricate networks of signaling molecules and transcriptional regulators^3,4^. Among these, the antagonistic action of classic phytohormones auxin (IAA) and cytokinin (CK) controls key aspects of meristematic activity including stem cell maintenance and the balance between proliferation and differentiation^2,5,6^. In the shoot apical meristem (SAM), CK signaling controls the size and positioning of the stem cells in the central zone, as well as cell cycle progression within the meristem^7–10^. By contrast, IAA signaling at the periphery of the SAM promotes organ initiation, thus differentiation^11–13^. The separation of hormone domains is fundamental for stem cell maintenance and is regulated by WUSCHEL – a key SAM regulator – which amplifies CK responses while attenuating IAA signaling regardless of the cellular auxin content^12^.

In the root apical meristem (RAM), the antagonistic action of IAA and CK specifies the distal stem cells (DSCs) during embryogenesis^14^. Upon the asymmetric division of the basal cell of the embryo root pole - the hypophysis - an auxin-mediated repression of CK signaling partitions hormone domains in the two daughters. As a result, CK signaling persists in the apical daughter where it specifies the stem cell organizer, the quiescent center (QC), while IAA signaling in the basal daughter specifies the columella stem cells (CSCs)^14^. During post-embryonic development, CSCs generate the outermost root tissue - the columella root cap - responsible for stem cell protection and gravity sensing^3^. Unlike other root stem cells, CSCs do not give rise to a transit-amplifying population but undergo a single asymmetric cell division (ACD)^3^. Following the ACD, the apical cells retain the CSC identity, while the basal daughters - the columella stem cell daughters (CSCDs) – briefly persist in an undifferentiated state before starting to accumulate starch granules as a marker of differentiation^3,15,16^. The initial partitioning of IAA and CK in root stem cells is later lost during post-embryonic development: CK signaling is switched off in DSCs, while an IAA signaling gradient spans across the distal tissues - from the QC to the outermost columella cell (CC) layer^15,17,18^.

The distal gradient of IAA is generated by a combination of local hormone biosynthesis and active auxin transport and controls all the aspects of the root cap development^15,19,20^. The IAA-mediated control of CSC fate is complex: while an auxin maximum promotes CSC division, an excess of the hormone causes their complete differentiation, leading to the division of the otherwise mitotically inactive QC to replenish the missing stem cells^3,15,18^. Specific components of the nuclear auxin signaling pathway (NAP), including repressive Class-C AUXIN RESPONSE FACTORs (ARF) and activating Class-A ARFs, were shown to regulate CSCs fate^18,21,22^. The auxin-mediated control of CSCs is generally thought to act in parallel with key stem cell regulators including WUSCHEL-LIKE HOMEOBOX 5 (WOX5), FEZ, SOMBRERO (SMB), and PLETHORA (PLT), although some points of contact among these pathways have been identified^16,18,23–25^. In particular, three Family II Homeodomain-Leucine Zipper (HD-Zip II) TFs were shown to act downstream of FEZ-borne proliferative input to counteract the auxin-induced differentiation of CSCs^24^.

Besides controlling the exit of proximal meristematic cells from the transit-amplifying population^26^, the functions of CK in DSC are still ill-defined. High CK levels were shown to interfere with the distal IAA maximum and to promote QC divisions via the typical transducers of the two-component signaling (TCS) pathway: the ARABIDOPSIS HISTIDINE KINASE (AHK) receptors, the positive Type-B ARABIDOPSIS RESPONSE REGULATORS (ARR) TFs, and their inhibitors Type-A ARR^27–30^.

Here, by studying the role of additional HD-Zip II family members in distal tissues, we uncover a positive role for CK in CSCs division. We show that CK and IAA antagonistically control DSCs fate through opposing pairs of cross-regulating HD-Zip II proteins. This regulatory circuit behaves as a bistable switch which precisely controls the balance between stemness and divisions in root distal tissues.

## RESULTS

### δ-and γ-clade HD-Zip II TFs oppositely regulate DSC divisions

The HD-Zip II TFs HAT3, ATHB4, and ATHB2 were shown to regulate CSC fate in Arabidopsis^24^. Phylogenetically, HAT3 and ATHB4 constitute the δ clade of HD-Zip II family, whereas ATHB2 with HAT1 and HAT2 form the sister clade γ^31^ (Fig. 1a). To understand whether HAT1 and HAT2 also regulate DSC fate, we analyzed the expression of GFP-tagged TFs (*pHAT1::HAT1-GFP* and *pHAT2::HAT2-GFP*, respectively). Within distal tissues, HAT1-GFP and HAT2-GFP were substantially overlapping with ATHB2-GFP, being all expressed in the QC, CSCs, and CSCDs, in opposition with δ-clade proteins which are usually excluded from DSCs^24^ (Fig.1b). Next, we generated *hat3 athb4 hat1* and *hat3 athb4 hat2* to analyze DSC phenotypes in comparison with *hat3 athb4* and *hat3 athb4 athb2*. In wild type (WT) plants the proportion of roots with one or two CSC layers is roughly 8:2, whereas *hat3 athb4* and *hat3 athb4 athb2* display an increment of differentiated CSCs (0 layer) at the expense of the two-layer pool, which is absent in the triple mutant^24^. Interestingly, *hat3 athb4 hat2* did not show any difference compared to *hat3 athb4*, whereas *hat3 athb4 hat1* displayed the most severe phenotype, reaching up to 40% of differentiated CSCs (Extended data Fig.1). While these results may suggest that δ-and γ-clade TFs redundantly control DSC fate, high-order γ-clade mutant combinations rather showed phenotypes opposite to *hat3 athb4*. Compared to the WT, *hat1 athb2* displayed a higher proportion of roots with 2 layers of CSCs coupled with increased QC divisions, whereas *hat1 athb2 hat2* displayed an increment of QC divisions but no CSC phenotypes (Fig.1c-e). Finally, *hat1 hat2* and *hat2 athb2* were similar to the WT (Fig.1d,e), indicating that only *hat1 athb2* mutant combinations promote DSC divisions. Next, we generated a β-estradiol-inducible version of *HAT1* (*XVE>HAT1)* to analyze the outcome of γ-gene overexpression on DSCs. Upon induction, *XVE>HAT1* displayed a dose-dependent increase in RAMs without CSCs (Fig.1f,g), the opposite of δ overexpressors^24^. Thus, HAT1 and ATHB2 (hereinafter the γ module) regulate DSC fate in opposition to HAT3 and ATHB4 (the δ module).

**Fig. 1.**
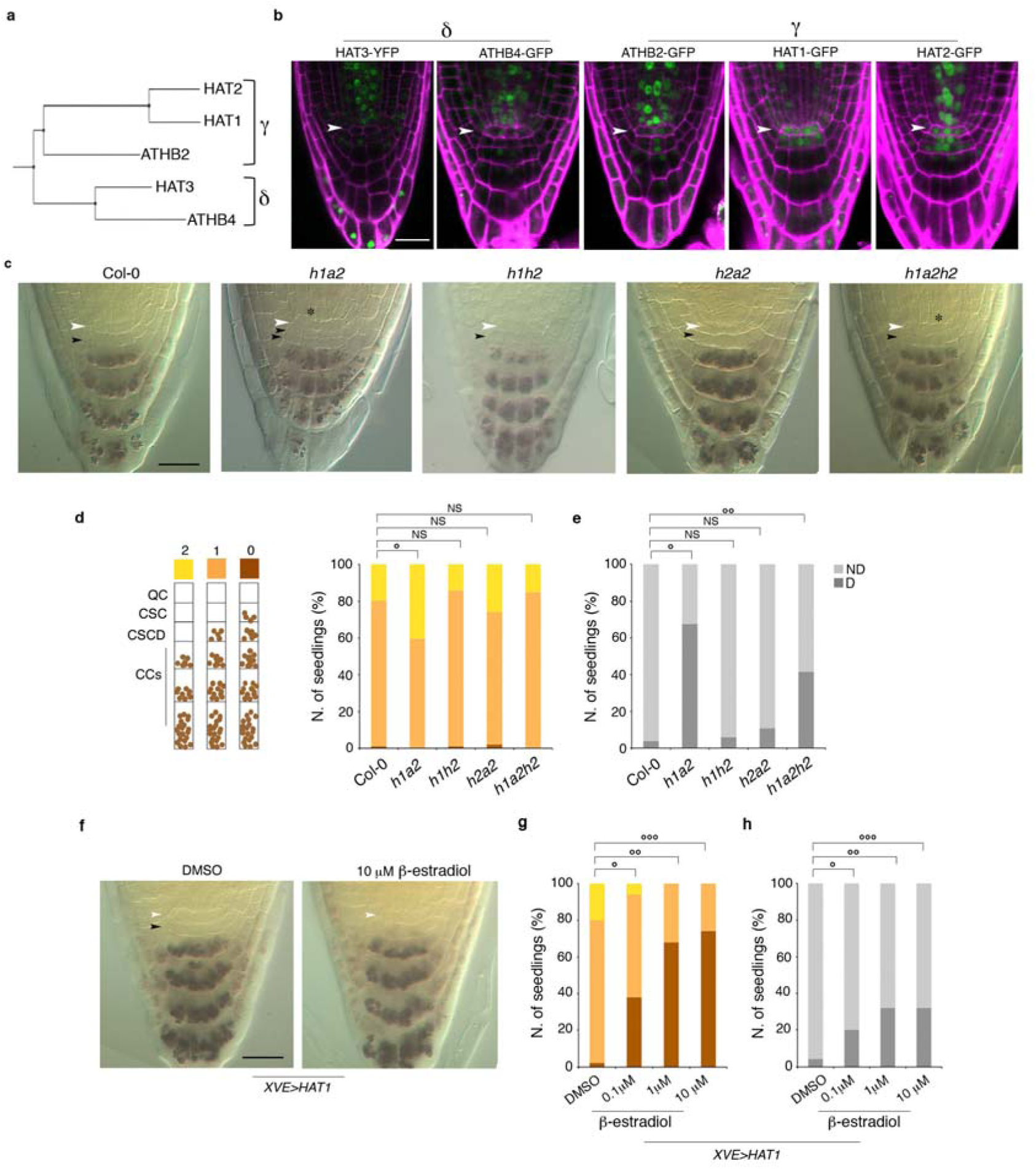
γ-clade HD-Zip II TFs negatively regulate DSC divisions. **a,** dendrogram of the five HD-Zip II TFs showing clades δ and γ. **b,** expression patterns of δ-clade (HAT3-YFP and ATHB4-GFP) and γ-clade (ATHB2-GFP, HAT1-GFP, and HAT2-GFP) HD-Zip II TFs in 5 day-old (d-o) arabidopsis roots counterstained with propidium iodide (PI, magenta), highlighting differences between δ (excluded from DSCs) and γ (expressed in DSCs) proteins. **c-e**, DSC phenotypes of 5d-o high order γ-clade mutant combinations showing enhanced CSC proliferation in *hat1 athb2* (*h1a2*) and QC divisions in *hat1 athb2* and *hat1 athb2 hat2* (*h1a2h2*). **c**, representative pictures of Lugol-stained distal tissues of WT (Col-0) and high order γ-clade mutants (*h1h2*, *hat1 hat2*; *h2a2*, *hat2 athb2*). Number of CSC layers (**d**) as exemplified by the scheme on the right and number of QC divisions (**e**) in the indicated genotypes (D, divided; ND, undivided). **d**, °*pVal_Col-0 vs h1a2_* = 0.00151, ^NS^*pVal* > 0.05; **e**, °*pVal_Col-0 vs h1a2_* = 1.47×10^-23^, °°*pVal_Col-0 vs h1a2h2_* = 2.36×10^-11^, ^NS^*pVal* > 0.05. **f-h,** DSC phenotypes of 5d-o *XVE>HAT1* seedlings treated for 24h with the indicated β-estradiol concentrations. Representative pictures of Lugol-stained *XVE>HAT1* roots (**f**), number of CSC layers (**g**), and number of QC divisions (**h**) in the indicated conditions. **g**, °*pVal_DMSO vs 0.1μM β-e_* = 1.49×10^-11^, °°*pVal_DMSO vs 1μM β-e_* = 4.52×10^-26^, °°°*pVal_DMSO vs 10μM β-e_* = 1.18 x10_-26_; **h**, °*pVal_DMSO vs 0.1μM β-e_* = 1.04×10^-4^, °°*pVal_DMSO vs1μM β-e_* = 8.35×10^-9^, °°°*pVal_DMSO vs 10μM β-e_* = 2.17 x10^-11^. White arrowheads, QC position; asterisks, QC divisions; black arrowheads, CSC layers. Scale bars: **b**,**c**,**f**, 20 μm.

**Fig. 2.**
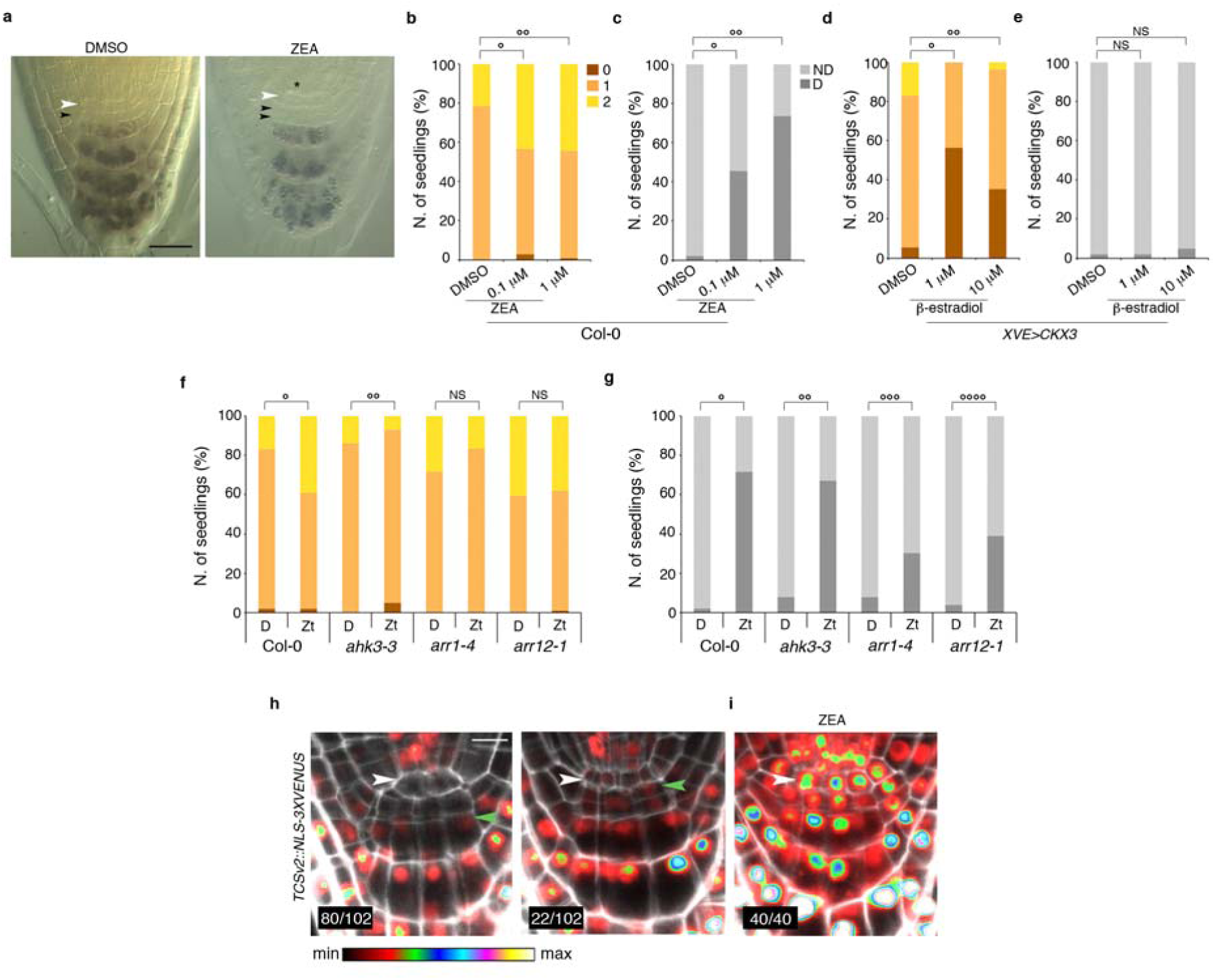
CK promotes CSCs division through canonical TCS pathway. **a-c,**CK treatments promote cell divisions in DSCs. **a**, representative images of DSCs from 5 d-o WT seedlings grown on 1μM ZEA or in control conditions (DMSO). Number of CSC layers (**b**) and QC divisions (**c**) in 5 d-o WT seedlings grown on increasing concentrations of ZEA or on DMSO as control. **b**, °*pVal_DMSO vs 0.1μM_* _ZEA_ = 4.41×10^-4^, °°*pVal_DMSO vs 1μM_* _ZEA_= 7.86×10^-4^; **c**, °*pVal _DMSO vs 0.1μM_* _ZEA_ = 1.24×10^-12^, °°*pVal_DMSO vs 1μM_* _ZEA_ = 2.99×10^-26^. White arrowheads, QC; asterisk, QC divisions; black arrowheads, CSC layers. **d-e**, reduction of endogenous CK via *XVE>CKX3* induction promotes CSC differentiation. Number of CSC layers (**d**) and QC divisions (**e**) in 5d-o *XVE>CKX3* seedlings upon induction with increasing β-estradiol concentrations for 24h. **d**, °*pVal_DMSO vs 1μM β-e_* = 2.78×10^-17^, °°*pVal_DMSO vs 10μM β-e_* = 9.25×10^-9^; **e**, ^NS^*pVal* > 0.05. **f-g**, Key components of CK signaling are required for CSC division. Number of CSC layers (**f**) and QC divisions (**g**) in Col-0, *ahk3-3*, *arr1-4*, and *arr12-1* seedlings grown for 5d with 1μM ZEA or in control conditions (DMSO). **f**, °*pVal _Col-0 D vs Col-0 Zt_* = 0.00088, °°*pVal _ahk3 D vs ahk3 Zt_* = 0.0170, ^NS^*pVal* > 0.05; **g**, °*pVal_Col-0 D vs Col-0 Zt_* = 3.07×10^-28^, °°*pVal _ahk3 D vs ahk3 Zt_* = 6.78×10^-19^, °°°*pVal _arr1 D vs arr1 Zt_* = 1.05×10^-4^, °°°°*pVal _arr12 D vs arr12 Zt_* = 4.26×10^-10^. **h-i**, *TCSv2::NLS-3xVENUS* expression in DSC correlates with divisions. In 5d-o WT seedlings (**h**), *TCSv2::NLS-3xVENUS* expression is excluded from CSC in ∼78% of samples (n=102) and present in CSC in the remaining ∼22% of root in association with cell division. In 5d-o WT seedlings grown on 1μM ZEA (**i**) *TCSv2::NLS-3xVENUS* expression peaks in dividing DSCs. TCSv2 intensity is depicted according to the false color scale (FCS) indicated below, PI is in white. White arrowheads, QC; green arrowheads, upper limit of TCSv2 expression in columella root cap. Scale bars: **a**, 20 μm; **h-i**, 10 μm.

**Fig. 3.**
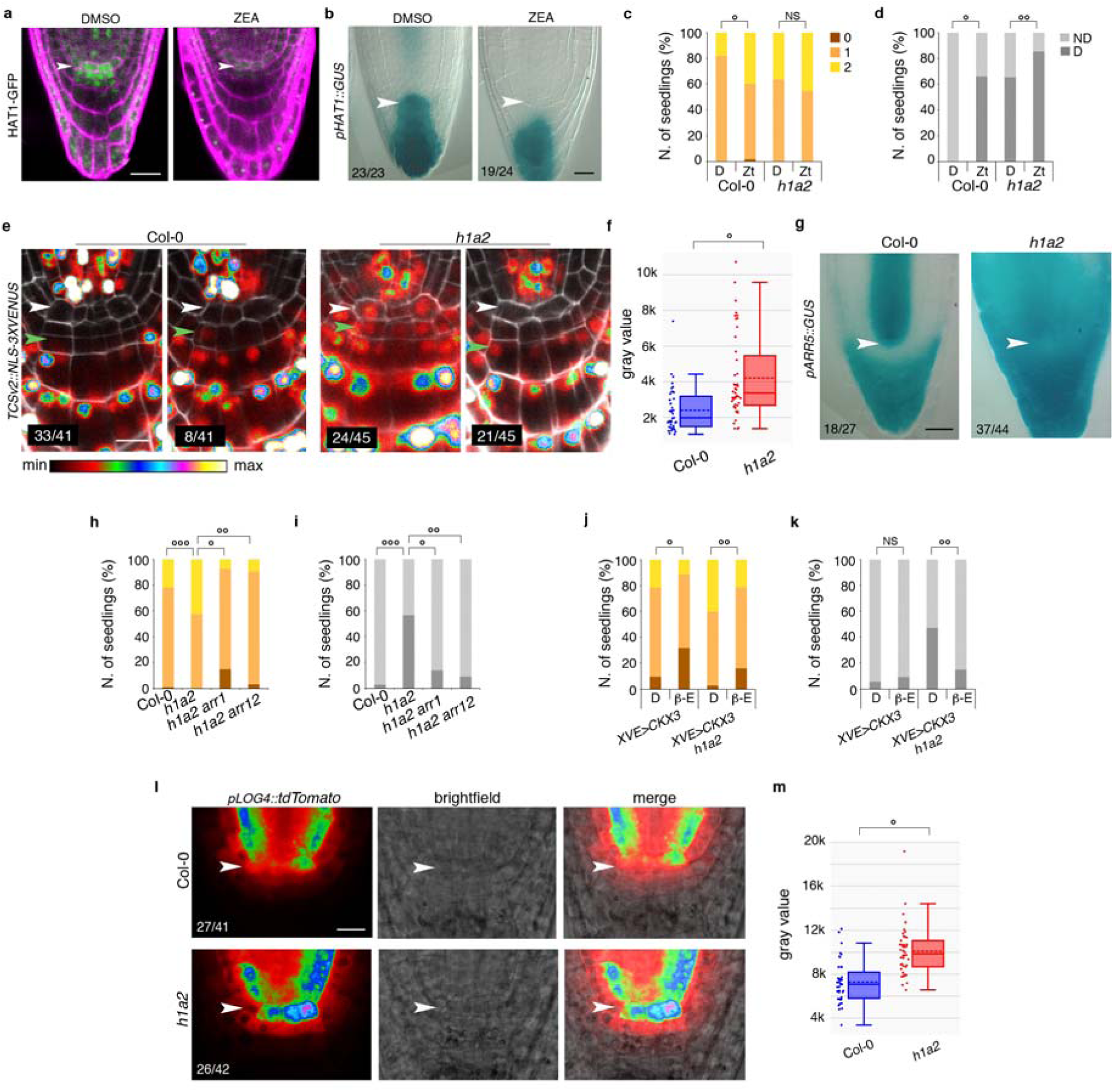
The γ module is repressed by CK and controls LOG4-mediated CK biosynthesis in DSCs. **a-b,**HAT1 is downregulated by CK in DSC. Expression of HAT1-GFP (**a**) and *pHAT1::GUS* (**b**) in 5 d-o roots grown in control conditions (DMSO) or on 0.1μM ZEA showing the disappearance of *HAT1* expression in ZEA-treated DSCs; white arrowheads, QC position **c-d**, reduced response of *hat1 athb2* mutant to CK. Number of CSC layers (**c**) and QC divisions (**d**) in 5d-o Col-0 and *hat1 athb2* (*h1a2*) seedlings grown with (Zt) or without (D) 1μM ZEA. **c**, °*pVal_Col-0 D vs Col-0 Zt_* = 9.96×10^-6^, ^NS^*pVal* > 0.05; **d**, °*pVal _Col-0 D vs Col-0 Zt_* = 1.12×10^-27^, °°*pVal_h1a2 D vs h1a2 Zt_* = 0.01027. **e-g**, increased CK signaling in *hat1 athb2*. **e**, representative images of *TCSv2::NLS-3xVENUS* expression in DSCs of 5d-o Col-0 and *h1a2* showing altered distribution of TCSv2 pattern in the double mutant compared to controls. White arrowhead, QC position; green arrowheads, upper limit of TCSv2 expression in columella. **f**, quantification of TCSv2 signal intensity in DSCs of seedlings grown as in (**e**), °*pVal_Col-0 Vs h1a2_ =* 2.86×10^-5^. **g**, representative images of *pARR5::GUS* expression in DSCs of 5 d-o Col-0 and *h1a2* roots; white arrowheads, QC position. **h-i**, Type-B ARRs are required for *h1a2* DSC phenotypes. Number of CSC layers (**h**) and QC divisions (**i**) in 5 d-o Col-0, *hat1 athb2*, *hat1 athb2 arr1*, and *hat1 athb2 arr12*, showing reversion of *hat1 athb2* DSCs phenotypes by the introgression of *arr* mutations. **h**, °*pVal_h1a2 vs h1a2arr1_* = 2.12×10^-9^, °°*pVal_h1a2 vs h1a2arr12_* = 1.24×10^-8^, °°°*pVal_Col-0 vs h1a2_* = 0.00177; **i**, *pVal_h1a2 vs h1a2arr1_* = 2.00×10^-6^, °°*pVal_h1a2 vs h1a2arr12_* = 9.16×10^-15^, °°°*pVal_Col-0 vs h1a2_* = 6.40×10^-19^. **j-k**, reduction of endogenous CK levels reverts *hat1 athb2* DSC phenotypes. Number of CSC layers (**j**) and QC divisions (**k**) in 5d-o *XVE>CKX3* and *XVE>CKX3 hat1 athb2* seedlings untreated (D) or treated for 24h with 10μM β-estradiol (β-E). **j**, °*pVal_XVE>CKX3 D vs β-e_* = 3.44×10^-4^, °°*pVal _XVE>CKX3 h1a2 D vs β-e_* = 2.45×10^-4^; **k**, ^NS^*pVal* > 0.05, °°*pVal _XVE>CKX3 h1a2 D vs β-e_* = 3.11×10^-7^. **l-m**, increased CK biosynthesis in DSCs of *hat1 athb2*. Representative images of *pLOG4::tdTomato* expression (**l**) in DSCs of 5 d-o Col-0 and *hat1 athb2*, visualized using the same FCS shown in (**e**); brightfield channel displays root anatomy. Notice increased *LOG4* expression in *hat1 athb2* QC (arrowheads) and CSCs compared to Col-0. **m**, quantification of *pLOG4::tdTomato* fluorescence in DSCs of plants grown as in (**l**), °*pVal_Col-0 vs h1a2_* = 2.86×10^-8^. Scale bars: **a**,**b**,**g**, 20 μm; **e**,**i**, 10 μm.

### CK promotes CSCs division via the TCS signaling pathway

The high number of QC divisions observed in γ-module mutants was reminiscent of plant grown on CK^28,30^. To understand whether CK promotes CSC divisions, we grew WT seedlings on 0.1μM and 1μM *trans*-Zeatin (ZEA) for 5 days. Compared to controls, ZEA treatments caused an increase in RAMs with 2 layers of CSCs in addition to QC divisions (Fig.2a-c). Interestingly, the increment in CSC layers already peaked at 0.1μM ZEA, whereas QC divisions displayed a dose-dependent increase (Fig.2b,c). Next, we used the *XVE>CKX3*^32^ line to reduce endogenous CK levels by overexpressing the catabolic gene *CYTOKININ OXIDASE 3* (*CKX3*). Following 24h β-estradiol treatments, we observed already at lower inducer concentrations a substantial increase in the number of RAMs with completely differentiated CSCs along with the disappearance of 2-layered meristems in comparison with mock-treated plants (Fig.2d,e). We next analyzed DSC phenotypes and CK responses in mutants of key transducers of CK signaling in the root: the AHK3 receptor, and the ARR1 and ARR12 Type-B regulators^26,30^. In *ahk3-3,* CK failed to promote CSC proliferation without affecting QC divisions, which are regulated by other receptors^30^ (Fig.2f,g). Similarly, *arr1-4* and *arr12-1* mutants did not display ZEA-induced CSCs proliferation while QC divisions were reduced compared with WT plants (Fig.2f,g). Thus, CK promotes DSC divisions through the canonical signaling pathway.

### CSC divisions are associated with local variations of distal CK signaling

To analyze the transcriptional readout of CK signaling in DSCs we used *TCSn::GFP*^33^ and *pARR5::GUS*^34,35^ markers. In WT plants, both markers are expressed in a gradient-like pattern across the columella, with a maximum in differentiated CCs and a minimum in DSCs (Extended data Fig.2a,b). Nevertheless, DSCs have the capacity to respond to CK as shown by the activation of *TCSn::GFP* and *pARR5::GUS* by short ZEA treatments (Extended data Fig.2a,b). This is consistent with the expression of the main Type-B ARRs - ARR1, ARR10, and ARR12 - in distal tissues (Extended data Fig.2c). To perform high-resolution analyses of CK signaling, we employed *TCSv2::NLS-3xVENUS*, a nuclear-localized marker with increased sensitivity^36^. TCSv2 expression was similar to TCSn, although we observed low-intensity signals across DSCs. In about 78% of samples (n=102), TCSv2 expression was excluded from the CSCs and QC; whereas in the remaining ∼22% of roots TCSv2 was expressed in CSCs with occasional signals in the QC (Fig.2h). Interestingly, TCSv2 expression in DSCs was usually associated with dividing cells (Extended data Fig.2d). Coherently, TCSv2 peaked in dividing DSCs of ZEA-grown roots (Fig.2i). These data indicate that DSC divisions are promoted by a local activation of CK signaling.

### HAT1 and ATHB2 are repressed by CK and regulate *LOG4* expression in DSCs

To explore the striking resemblance between *hat1 athb2* and CK-treated plants we analyzed *pATHB2::ATHB2-GFP* and *pHAT1::HAT1-GFP* expression upon 5d treatments with ZEA. Both HAT1-GFP and ATHB2-GFP were repressed by CK likely at the transcriptional level as we observed *pHAT1::GUS* expression disappearing from DSCs upon prolonged ZEA treatments (Fig.3a,b; Extended data Fig.3a-c). Interestingly, *hat1 athb2* CSCs did not respond to ZEA treatments, whereas the QC only displayed a moderate increment in divisions compared to the control (Fig.3c,d), suggesting high CK activity. Thus, we introgressed *pARR5::GUS* and *TCSv2::NLS3xVENUS* in the double mutant. Both markers displayed stronger expression in *hat1 athb2* distal tissues compared to the WT (Fig.3e-g). Also, we observed a higher proportion of TCSv2-expressing DSCs in *hat1 athb2* (53.3%; n=45) compared to the WT (19.5%; n=41), coherently with mutants’ higher division rates (Fig.3e). We next generated *hat1 athb2 arr1* and *hat1 athb2 arr12* mutants and found that the introgression of either *arr* mutation reverted the *hat1 athb2* DSC phenotypes to WT-like levels (Fig.3h,i). However, the expression of Type-B ARRs in *hat1 athb2* was similar to the WT (Extended data Fig.3f-i), showing that CK signaling activation is not associated with altered expression of ARR1 and ARR10. Thus, we introgressed *XVE>CKX3* in *hat1 athb2* to reduce endogenous CK content. Upon β-estradiol treatments, *XVE>CKX3 hat1 athb2* plants displayed a strongly reduced number of CSC layers and QC division compared to controls (Fig.3j,k), indicating that *hat1 athb2* DSC phenotypes are caused by high CK levels. We focused on LONELY GUY 4 (LOG4), a rate-limiting biosynthetic enzyme involved in stem cell patterning^7^. Relevantly, *pLOG4::tdTomato*^37^ expression was higher in *hat1 athb2* DSCs than in the WT (Fig.3l,m), suggesting that the γ module regulates DSC divisions through LOG4-derived CK.

### HAT1 and ATHB2 regulate DSC fate downstream of ARF8-mediated auxin signaling

γ proteins were previously linked to IAA^24,38^, therefore we introgressed *DR5::VENUS-N7* in *hat1 athb2* and observed a substantial reduction in DR5 expression, similar to CK-treated WT plants^29,39^ (Extended data Fig.4). To understand whether high CK affects auxin responses in DSCs, we carried out NAA treatments on ZEA-conditioned WT plants. NAA promoted CSC differentiation to the same extent in CK-conditioned and control plants alike (Fig.4a-c). However, NAA-induced CSC differentiation was impaired in *hat1 athb2* (Fig.4d-f), suggesting that auxin counteracts CK responses in DSCs through the γ module. Indeed, ATHB2 is upregulated by auxin^24^, and *pHAT1::GUS* expression displayed a substantial upregulation in time course analyses of NAA response, peaking at 24h time-point (Fig.4g; Extended data Fig.5a). We next used *XVE>HAT1* to uncouple HAT1 expression from hormone-mediated regulation and carried out β-estradiol treatment on CK-conditioned plants. Interestingly, β-estradiol caused CSC differentiation in ZEA-conditioned and control plants alike (Fig.4h,i), demonstrating that - like auxin - HAT1 represses CK-mediated DSC divisions.

**Fig. 4.**
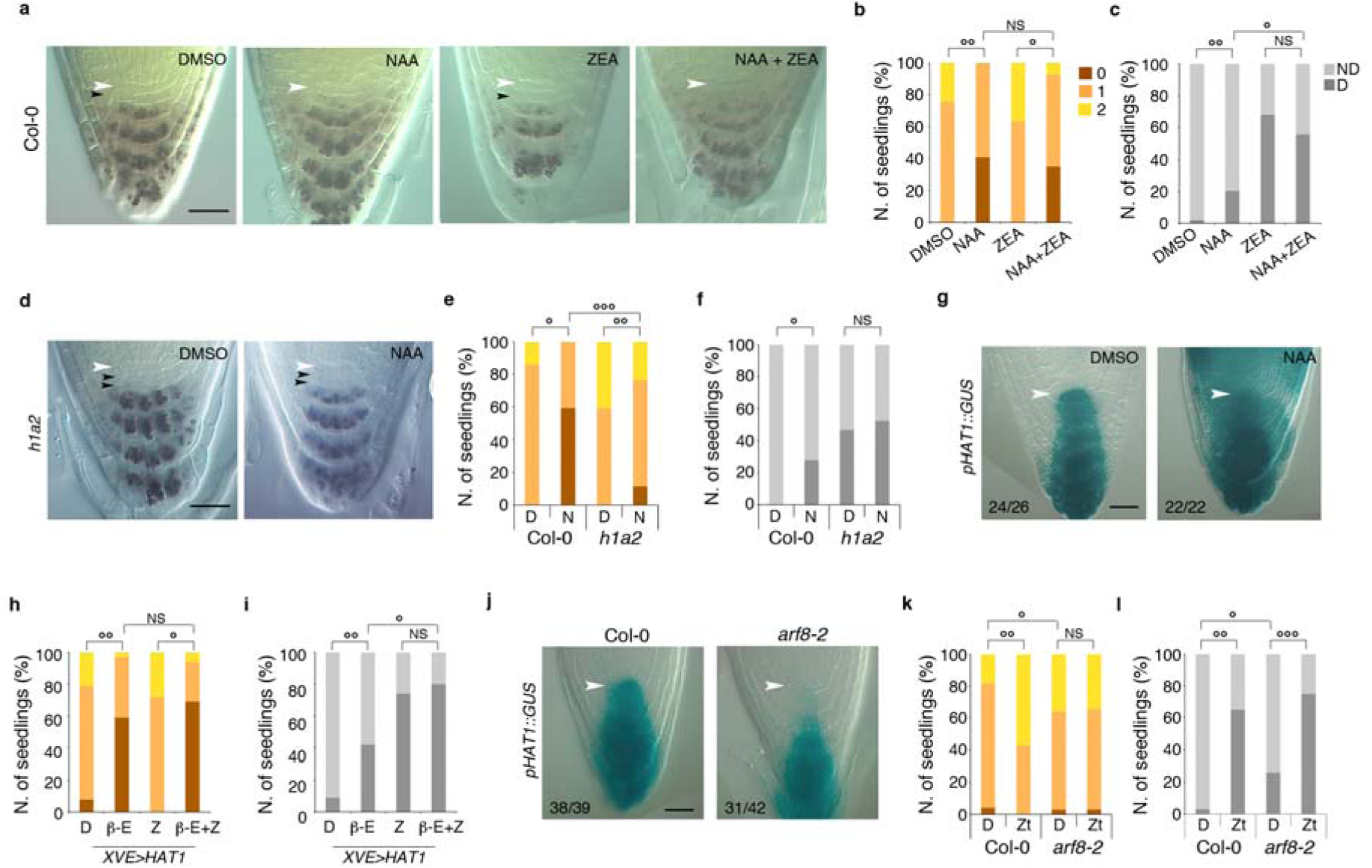
IAA counteracts CK-mediated DSC divisions via the γ module. **a-c,** NAA overrides CK-mediated CSC proliferation. Representative images (**a**), number of CSC layers (**b**), and QC divisions (**c**) in Col-0 roots grown for 4d on 1μM ZEA and then treated for 24h with 1μM ZEA alone or 1μM ZEA/5μM NAA. 5d DMSO and 4d DMSO with 24h 5μM NAA were used as controls. NAA completely overrides CK-mediated CSC proliferation (**a,b**) but has no effect on QC divisions (**a,c**). **b**, °*pVal_ZEA vs NAA+ZEA_* = 3.89×10^-18^, °°*pVal_DMSO vs NAA_* = 1.18×10^-19^, ^NS^*pVal* > 0.05; **c**, °*pVal_NAA vs NAA+ZEA_* = 1.19×10^-8^, °°*pVal_DMSO vs NAA_* = 5.42×10^-7^, ^NS^*pVal* > 0.05. White arrowheads, QC position; black arrowheads CSC layers. **d-f**, *hat1 athb2* DSCs have impaired responses to NAA. Representative images (**d**), number of CSC layers (**e**), and QC divisions (**f**) in 5 d-o *hat1 athb2* roots untreated (D) or treated with 5μM NAA (N). **e**, °*pVal_Col-0 D Vs Col-0 N_* = 8.32×10^-20^, °°*pVal_h1a2 D Vs h1a2 N_*= 3.47×10^-4^, °°°*pVal_h1a2 N vs Col-0 N_*= 8.42×10^-14^; **f**, °*pVal_Col-0 D Vs Col-0 N_* = 5.42×10^-7^, ^NS^*pVal* > 0.05. White arrowheads, QC position; black arrowheads CSC layers. **g**, *HAT1* is strongly induced by auxin. Representative images of *pHAT1::GUS* roots untreated (DMSO) or treated for 24h with 5μM NAA; arrowheads, QC position. **h-i**, HAT1 overexpression overrides CK-induced proliferation. Number of CSC layers (**h**), and QC divisions (**i**) in *XVE>HAT1* roots grown for 4d on 1μM ZEA and then treated for 24h with 1μM ZEA alone (Z) or 1μM ZEA/10μM β-estradiol (β-E+Z). 5d DMSO (D) and 4d DMSO with 24h 10μM β-estradiol (β-E) were used as controls. **h**, °*pVal _Z vs β-e+Z_* = 2.61×10^-29^, °°*pVal _D vs β-e_* = 1.01×10^-15^, ^NS^*pVal* > 0.05; **i**, °*pVal _β-e vs β-e+Z_* = 5.19×10^-8^, °°*pVal _D vs β-e_* = 9.25×10^-8^, ^NS^*pVal* > 0.05. **j**, ARF8 drives *HAT1* expression in DSCs. Representative images of *pHAT1::GUS* expression in 5 d-o Col-0 and *arf8-2*, showing the disappearance of *HAT1* expression in mutants’ DSCs; arrowheads, QC position. **k-l**, *arf8-2* displays DSCs phenotypes and CK insensitivity like *hat1 athb2.* Number of CSC layers (**k**), and QC divisions (**l**) in 5 d-o Col-0 and *arf8-2* roots untreated (D) or treated with 1μM ZEA (Zt). **k**, °*pVal_Col-0 D vs arf8 D_* = 0.01643, °°*pVal_Col-0 D vs Col-0 Zt_* = 9.25×10^-9^, ^NS^*pVal* > 0.05; **l**, °*pVal_Col-0 D vs arf8 D_* = 3.43×10^-6^, °°*pVal_Col-0 D vs Col-0 Zt_* = 1.27×10^-22^; °°°*pVal_arf8 D vs arf8 Zt_* = 7.20×10^-12^. Scale bars: **a**,**d**,**g**,**j**, 20 μm.

**Fig. 5.**
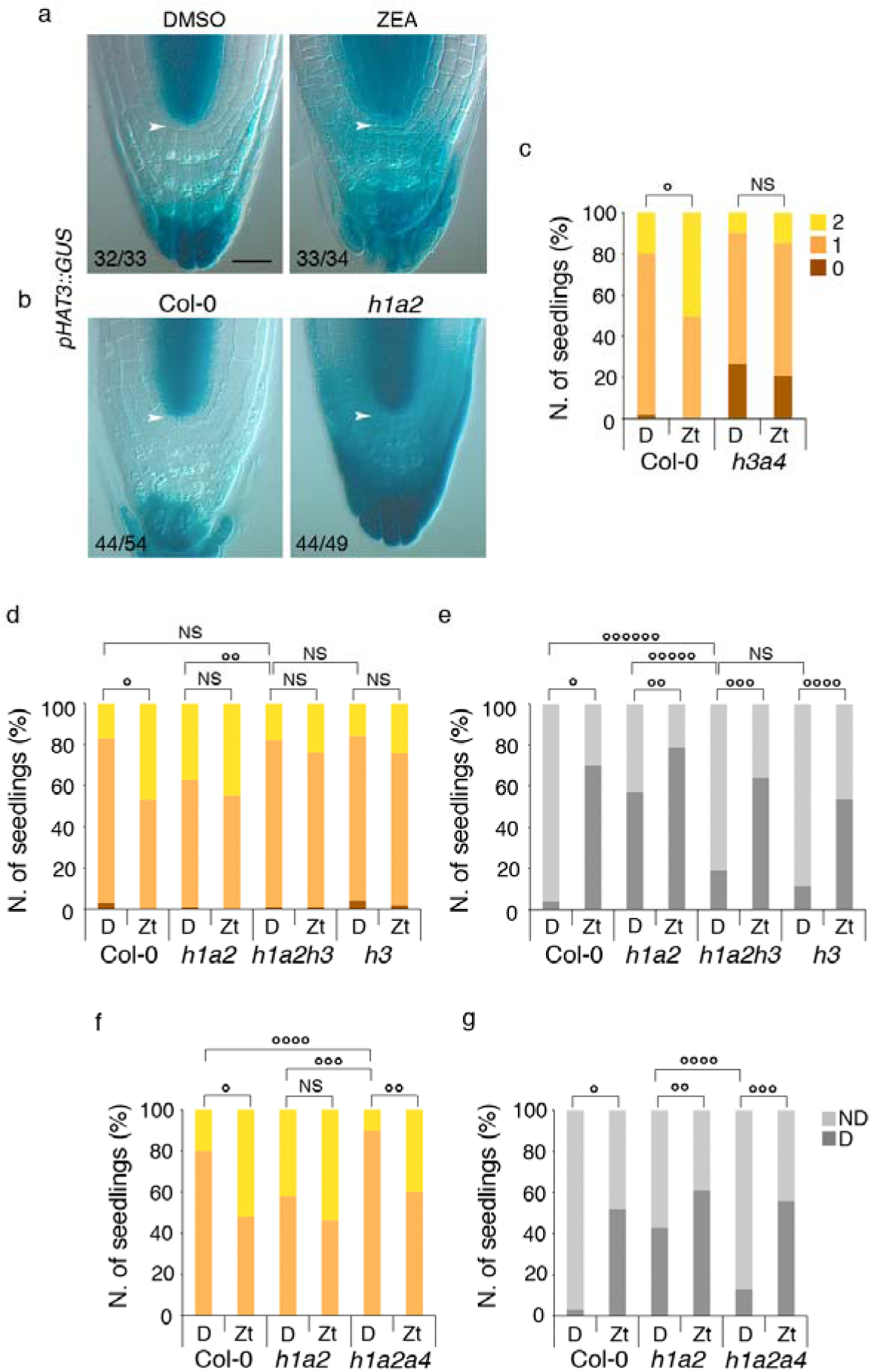
The γ module promotes DSC division downstream of CK and the γ module. **a-b,** *HAT3* is upregulated in DSCs by prolonged CK-treatments. Representative images of *pHAT3::GUS* expression in DSCs of roots treated with 1μM ZEA for 5d compared to untreated (DMSO) controls (**a**), and in DSCs of *hat1 athb2* (*h1a2*) compared to Col-0 (**b**). Arrowheads, QC position. **c**, *hat3 athb4* DSCs do not respond to CK. Number of CSC layers in 5d-o Col-0 and *hat3 athb4* (*h3a4*) plants untreated (D) or treated with 1μM ZEA (Zt); °*pVal_Col-0 D Vs Col-0 Zt_* = 3.74×10^-6^, ^NS^*pVal* > 0.05. **d-e**, HAT3 is required for CK-induced DSC divisions. Number of CSC layers (**d**), and QC divisions (**e**) in 5 d-o Col-0, *h1a2*, *hat1 athb2 hat3* (*h1a2h3*), and *hat3* (*h3*) roots untreated (D) or treated with 1μM ZEA (Zt), showing that *h3* mutation impairs divisions in CSC, and partly in QC, downstream of CK and the γ module. **d**, °*pVal_Col-0 D vs Col-0 Zt_* = 2.95×10^-6^, °°*pVal_h1a2 D vs h1a2h3 D_* = 0.00302, ^NS^*pVal* > 0.05; **e**, °*pVal_Col-0 D vs Col-0 Zt_* = 2.33×10^-24^, °°*pVal_h1a2 D vs h1a2 Zt_* = 8.12×10^-4^, °°°*pVal_h1a2h3 D vs h1a2h3 Zt_* = 1.19×10^-24^, °°°°*pVal_h3 D vs h3 Zt_* = 3.50×10^-10^, °°°°°*pVal _h1a2 D vs h1a2h3 D_* = 1.74×10^-8^, °°°°°°*pVal _Col-0 D vs h1a2h3 D_* = 0.00141, ^NS^*pVal* > 0.05. **f-g**, ATHB4 is required for DSC divisions downstream of the γ module. Number of CSC layers (**f**), and QC divisions (**g**) in 5 d-o Col-0, *h1a2*, and *hat1 athb2 athb4* (*h1a2a4*) roots untreated (D) or treated with 1μM ZEA (Zt), showing that the introgression of *a4* mutation restores WT-like phenotypes and CK responses in *h1a2*. **f**, °*pVal_Col-0 D vs Col-0 Zt_* = 6.08×10^-5^, °°*pVal_h1a2a4 D vs h1a2a4 Zt_* = 9.20×10^-12^, °°°*pVal_h1a2 D vs h1a2a4 D_* = 5.64×10^-10^, °°°°*pVal_Col-0 D vs h1a2a4 D_* = 0.00451, ^NS^*pVal* > 0.05; **g**, °*pVal_Col-0 D vs Col-0 Zt_* = 4.02×10^-16^, °°*pVal_h1a2 D vs h1a2 Zt_* = 0.01591, °°°*pVal_h1a2a4 D vs h1a2a4 Zt_* = 1.45×10^-10^, °°°° *pVal_h1a2 D vs h1a2a4 D_* =3.33×10^-6^. Scale bar: **a**, 20 μm.

**Fig. 6.**
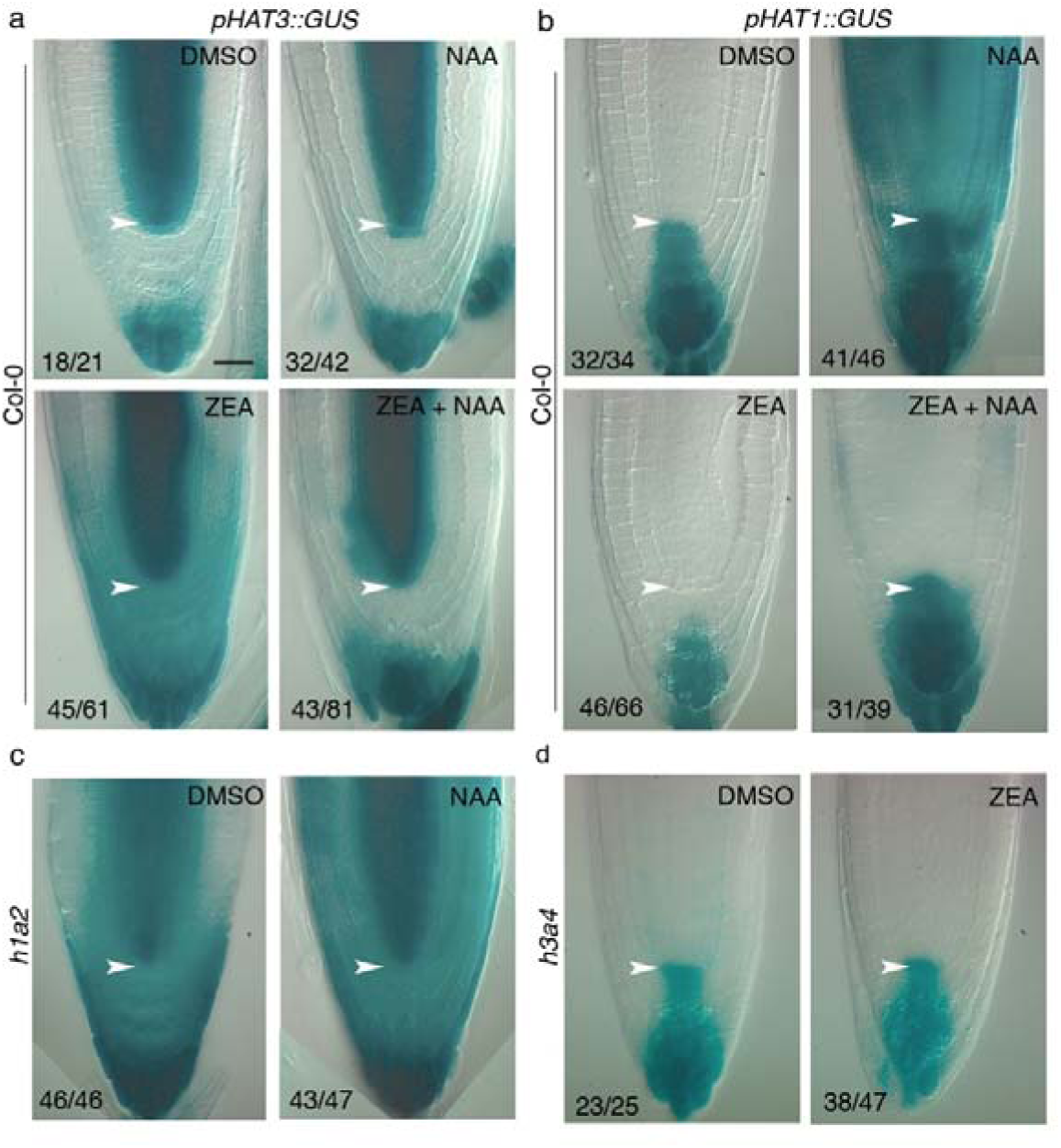
Cross-regulation between δ and γ modules downstream of hormones. **a-b,** Representative images of 5 d-o *pHAT3::GUS* (**a**) and *pHAT1::GUS* (**b**) roots in ZEA/NAA co-treatments. Upper panels: plants were grown for 4d on DMSO and either kept on DMSO or treated with 5μM NAA (NAA) for 24h. Lower panels: plants were grown for 4d on 1μM ZEA and then treated for 24h with 1μM ZEA alone (ZEA) or 1μM ZEA/5μM NAA (ZEA+NAA). Notice that NAA represses ZEA-induced *HAT3* expression in DSC but has no effect in absence of ZEA (**a**). By contrast, NAA restores *HAT1* expression in DSC overcoming ZEA-mediated downregulation (**b**). **c,** the γ module represses *HAT3* expression in DSCs downstream of auxin. Representative images of 5 d-o *hat1 athb2 pHAT3::GUS* roots untreated (DMSO) or treated for 24h with 5μM NAA showing lack of NAA-mediated downregulation of *HAT3* in DSCs. **d,** the δ module represses *HAT1* expression downstream of CK. Representative images of 5 d-o *hat3 athb4 pHAT1::GUS* roots untreated (DMSO) or treated for 5d with 1μM ZEA showing lack of CK-mediated downregulation of *HAT1* in DSCs. Arrowheads, QC position. Scale bar: 20 μm.

We observed that the auxin-mediated induction of *pHAT1::GUS* was blocked by auxinole, a competitive inhibitor of the TIR1/AFB-Aux/IAA co-receptor complex^40^ (Extended data Fig.5b), suggesting that *HAT1* expression depends on Class-A ARFs activity. Among these, only ARF8 is specifically expressed in distal tissues^41^, thus we introgressed *pHAT1::GUS* in *arf8-2* and found that *HAT1* was not expressed in mutants’ DSCs (Fig.4j). Coherently, *arf8-2* displayed an increased number of CSC layers and QC divisions compared to the WT, and was largely impaired in distal responses to ZEA, like *hat1 athb2* (Fig.4k,l). Taken together, our data demonstrate that auxin regulates γ module expression via ARF8 to repress CK-induced DSC divisions.

### HAT3 and ATHB4 promote CSC divisions downstream of CK signaling

The δ module promotes CSC divisions in opposition to IAA^24^ and there is evidence that Type-B ARRs bind the promoters of *HAT3* and *ATHB4* upon prolonged CK stimulus^8^ (Extended data Fig.6a,b), indicating that these TFs might act downstream of CK. Both *pHAT3::GUS* and *pATHB4::GUS* were induced by 5d ZEA treatments, although *pHAT3::GUS* expression was specifically upregulated in DSCs (Fig.5a; Extended data Fig.6c). Coherently, *pHAT3::GUS* was strongly upregulated in *hat1 athb2* DSCs compared to the WT (Fig.5b). The upregulation of *pHAT3::GUS* in *hat1 athb2* was largely abolished by the introgression of *arr12* (Extended data Fig.6d), indicating that the δ module acts as CK effector in DSCs. In fact, when grown on ZEA for 5d *hat3 athb4* did not display increased CSC divisions (Fig.5c) confirming that CK-induced proliferation requires δ proteins. We next generated the triple mutants *hat1 athb2 hat3* and *hat1 athb2 athb4* and observed that the introgression of either *δ* mutation reverted the double mutant CSC phenotype and substantially reduced the number of QC divisions (Fig.5d-g). When grown on ZEA, *hat1 athb2 athb4* displayed WT-like DSC responses, whereas *hat1 athb2 hat3* was impaired in CSC proliferation, but not QC divisions, like the single *hat3* mutant (Fig.5d-g). These data demonstrate that δ proteins regulate DSCs divisions downstream of CK signaling, with HAT3 playing a dominant role.

### Transcriptional cross-regulation between γ and δ modules controls DSC fate downstream of hormones

The cross-regulation of HD-Zip II TFs is well established^31,42–44^, thus we figured that it may occur in DSCs. Therefore, we analyzed *pHAT3::GUS* and *pHAT1::GUS* expression in ZEA/NAA cotreatments. NAA alone did not affect *pHAT3::GUS* pattern, however in ZEA-conditioned plants NAA repressed the CK-driven *HAT3* expression in DSCs (Fig.6a). Conversely, in ZEA-conditioned plants NAA restored *pHAT1::GUS* expression in DSCs, overriding the CK-mediated downregulation (Fig.6b). Interestingly, NAA failed to downregulate *pHAT3::GUS* expression in *hat1 athb2* (Fig.6c), demonstrating that the γ module represses *HAT3* - hence DSC divisions - downstream of auxin. Next, we analyzed *pHAT1::GUS* expression in *hat3 athb4* and found no difference compared to the WT (Fig.6d). However, ZEA failed to downregulate *pHAT1::GUS* expression in *hat3 athb4* DSCs (Fig.6d), demonstrating that the δ module regulates *HAT1* downstream of CK. Our data demonstrate that IAA and CK regulate DSC fate through opposing pairs of HD-Zip II TFs.

## DISCUSSION

In this work, by studying the complex genetic relationships of HD-Zip II TFs in root stem cells, we identified CK as a novel proliferative input that controls DSC fate. CK promotes DSC division via δ proteins and its action is counteracted by IAA via the γ module. In this framework, the cross-regulation of HD-Zip II TFs plays a central role in translating hormone-borne information into cell fate decisions. The mutual inhibition of TFs downstream of antiparallel gradients of diffusible molecules is typical of the bistable switch, a regulatory network topology used by cells to translate graded morphogen inputs into sharp developmental outputs like discrete gene expression domains^45,46^. In our model, the IAA maximum drives the expression of the γ module in DSCs via ARF8, leading to the local downregulation of δ proteins and CK synthesis. As a result, the δ module is confined in underlying cells, sustained by columella-specific CK signaling. This configuration inhibits DSC divisions and constitutes the stable state 1. Upon CK signaling activation, the long-term upregulation of the δ module by Type-B ARRs slowly represses γ module activity, resulting in a local increase of LOG4-derived CK and a further induction of δ proteins in DSCs. This promotes DSC division and constitutes the stable state 2 (Extended data Fig.7). Bistable switches are characterized by hysteresis, i.e. the memory of a predefined stable state as a means to buffer against fluctuations in gene expression^45,46^. Here, the different hormone response dynamics may contribute to hysteresis: the γ module is activated by auxin within hours; the δ module is activated by CK within days, coherently with the timing of CSC divisions in Arabidopsis roots^47^. Thus, the promptness of γ-module response to IAA might help restore stable state 1, overriding the slow-operating CK-δ branch. This is further supported by previous evidence of auxin signaling pulses occurring in-between CSC division events^15^, suggesting that the IAA-γ branch may reinstate CSC stemness and promote CSCD differentiation at once. Additional feedback regulations may incorporate into this circuit: HAT1 was shown to potentiate IAA signaling^38^, while the δ module alters IAA transport and Class-C ARF expression^24^. Noteworthy, QC divisions, while being controlled by the γ module via LOG4-mediated CK synthesis, are only partly regulated by δ proteins and mostly dependent on Type-B ARRs as shown by previous works^30^. Also, the role of HAT2 within the network remains ill-defined. However, our genetic analyses suggest that HAT2 function in CSCs may align with that of δ proteins, rather than of γ-clade paralogs.

The role of CK in root stem cell division is coherent with previous observations^28,30^, albeit in contrast with the evidence of a CK-mediated promotion of RAM differentiation^26^. This apparent contradiction rather suggests a context-dependent role of CK in stem cell fate control. In this perspective, local auxin levels might change the outcome of CK action. For instance, in the SAM auxin levels are high and CK promotes proliferation^9,12^, whereas in the RAM transition zone IAA levels are at a minimum and CK promotes differentiation^26,48^. In DSCs, high IAA levels may confer CK a proliferative role, while converging antiparallel IAA and CK signals regulate the precise switch from stemness to proliferation and back via HD-Zip II TFs.

In conclusion, our work identifies CK as a key player in DSC fate regulation and expands the current knowledge of IAA/CK antagonism in meristematic tissues. It remains to be established how the HD-Zip II switch wires up to other DSC regulatory circuits. Previous evidence suggests that both δ and γ modules may act downstream of FEZ^24^, however interactions with other key regulators of DSC fate including WOX5 – PLTs complexes^49,50^ will need to be explored in future works.

## MATERIALS AND METHODS

### Plant Material and growth conditions

Plants were grown vertically for 5 days as previously described^24^. HD-Zip II lines used for this study were previously described: *hat3-3*, *athb4-1*, *hat3-3 athb4-1*, *hat3-3 athb4-1 athb2-3*, *pHAT3::GUS*, *pATHB4::GUS*^44^; *pHAT3::HAT3-YFP*; *pATHB4::ATHB4-GFP*, *pATHB2::ATHB2-GFP*^24^; *hat1-2*, *hat2-1*^51^. Other mutants and marker lines used in this study were previously described: *arr1-4*, *arr12-1*^52^; *ahk3-3*^53^; *arf8-2*^54^; *XVE>CKX3*^32^; *DR5::VENUS-N7*^11^; *TCSn::GFP*^33^; *TCSv2::NLS-3xVENUS*^36^; *pARR1::ARR1-Ypet*, *pARR10::ARR10-Ypet, pARR12::ARR12-Ypet*^8^; *pARR5::GUS*^35^; *pLOG4::tdTomato*^37^.

High order mutant combinations used in this study were generated by crossing: *hat3-3 athb4-1 hat1-2*, *hat3-3 athb4-1 hat2-1*, *hat1-2 athb2-3*, *hat1-2 athb2-3 hat2-1*, *hat1-2 hat2-1*, *athb2-3 hat2-1*, *hat1-2 athb2-3 hat3-3, hat1-2 athb2-3 athb4-1, hat1-2 athb2-3 arr1-4, hat1-2 athb2-3 arr12-1.* All the multiple mutants were selected in F2 by PCR genotyping and re-analyzed in F3 (see Extended Data Table1 for primer details). Marker lines were introgressed in single or double mutants by crossing and the progenies were screened in F2 by phenotyping and/or PCR genotyping. Homozygosity of all the reporters was determined either by resistance selection or by unanimous GUS/Fluorescent Protein signals in F3 (n≥30). For all the crosses, control lines for mutants and markers were resegregated from the F2 progeny.

The *XVE>HAT1* construct was generated by inserting the *HAT1* cDNA in pER8 vector^55^ by restriction/ligation cloning. The *pHAT1::GUS* was generated by inserting a promoter fragment of 2Kb upstream of the *HAT1* start codon in the pMDC163 vector^56^ via Gateway cloning. *pHAT1::HAT1-GFP* and *pHAT2::HAT2-GFP* constructs were generated by inserting DNA fragments encompassing 2Kb upstream of the start codon and the entire coding sequence of *HAT1* and *HAT2*, respectively, in the pMDC107 vector^56^ by Gateway cloning. Details and sequences of cloning primers are given in Extended Data Table1). Constructs were transformed in Col-0 plants via simplified floral dip procedure^57^. Transformants were identified by resistance marker selection, and experiments were carried out on homozygous T3 seedlings.

### Chemical treatments

Long-term treatments were performed by growing plants directly on a medium supplemented with the corresponding chemical for 5 d: ZEA concentrations, 0.1μM or 1 μM. Short-term treatments were carried out by transferring 4-5 d-o seedlings onto plates supplemented with the corresponding chemical and further incubated for the required times: NAA, 5μM for 1h, 4h, 24h; β-est, 0.1μM, 1μM, and 10μM for 24h. Equal amounts of DMSO were used as negative controls for all the treatments. ZEA/NAA cotreatments were performed by growing plants on 1μM ZEA for 4d and seedlings were subsequently transferred onto plates containing either 1μM ZEA alone or 1μM ZEA/5μM NAA for additional 24h. Control plants were grown on DMSO and then either transferred on 5μM NAA or kept on DMSO for additional 24h. ZEA/β-est cotreatments were performed using an identical experimental setup with the following concentrations: 1μM ZEA; 10μM β-est, and 1μM ZEA/10μM β-est. NAA/auxinole cotreatments were performed by transferring 5d-o seedlings on plates containing either 5μM NAA or 5μM NAA/10μM auxinole for 4h.

### Phenotypic analyses

DSC analysis was carried out on Lugol-stained root apices as previously described^24^. The distribution of CSC layers was determined as follows: 2 layers, if at least 3 CSCs on the middle focal plane were divided and CSCD did not display Lugol-stained starch granules; 1 layer, if CSCs were undivided or if at least 1 CSCD displayed Lugol-stained starch granules; 0 layer; if at least 1 CSC displayed Lugol-stained starch granules. QC divisions were assessed by scoring the presence of a complete periclinal cell wall in QC cells imaged on the middle focal plane. All the experiments were carried out at least twice to ensure reproducibility, and a minimum of 70 samples were analyzed for each genotype/condition.

### Microscopy and histochemistry

β-glucuronidase staining experiments were carried out substantially as previously described^58^. Staining times were adjusted for each marker/condition used: *pHAT3::GUS* and *pATHB4::GUS*, 2h staining; *pHAT1::GUS*, 1h30’ staining; *pARR5::GUS*, 2h staining in *hat1 athb2* experiments, 30’ staining in CK treatments. Expression patterns were observed on chloral hydrate-cleared samples under a Zeiss Axioskop 2 plus microscope equipped with Nomarski optics as previously described^24^. Confocal microscopy imaging was carried out using a Zeiss LSM 700 as previously described^24^. Quantifications of fluorescent signal intensity were carried out using Fiji (https://imagej.net/software/fiji/). For signal intensity quantifications in the DSCs, freehand selection tool was used in the PI channel to draw a region of interest (ROI) encircling the DSCs; ROI was used to measure the signal intensity in the Fluorescent Protein channel. Each experiment was repeated at least twice to ensure reproducibility and a minimum of 20 samples were analyzed for each line/condition.

## Statistical analyses

Statistical analyses of the distribution of QC divisions and CSC layers were carried out in VassarStats - Website for Statistical Computation (http://vassarstats.net/index.html) using: for the QC, 2×2 contingency tables followed by two-tailed Fisher’s exact probability test; for CSCs, 2×3 contingency tables followed by two-tailed Fisher-Freeman-Halton exact probability test. Statistical analyses of fluorescent protein intensity were carried out in MS Excel by using a two-tailed Student’s *t*-test. Statistical analyses of fluorescent protein distribution in distal tissues were carried out in MS Excel by using a *χ^2^* test.

### Phylogenetic Analysis

HD-Zip II alignment was obtained in InParanoiDB 9 database (https://inparanoidb.sbc.su.se/) based on the N-ter domain (*HD-ZIP_N;* Pfam, PF04618) homology. The tree was subsequently visualized in phylo.io (https://phylo.io/viewer/). The orthologous outgroup was removed for clarity.

## Acknowledgements

We would like to thank Bert de Rybel and Maya Bar for generously sharing seeds; Giorgio Morelli and Jakub Hajny for discussions and critical reading of the manuscript. Work in authors’ laboratory is funded by Agritech National Research Center within the European Union Next Generation EU program (the National Recovery and Resilience Plan, mission 4, component 2, investment 1.4 D.D. 1032 17/06/2022, project CN00000022, CUP-B83C22002840001), and by the Regione Lazio POR-FESR 2014-2020 (SMARTBREED project - B85F21003370002).

## Author contribution

G.S. and M.S. designed the research, defined the methodology, and supervised the project.

G.S., M.S., M.P., I. B., G.Sa., and A.A. conducted the experiments. A.D.P. generated genetic resources. G.S. created the visualization. M.S. wrote the manuscript with inputs from other authors.

