## Supplementary Information for "Auxin and cytokinin antagonistically regulate Arabidopsis root distal stem cell fate through opposing pairs of HD-Zip II transcription factors"

### 1 EXTENDED DATA

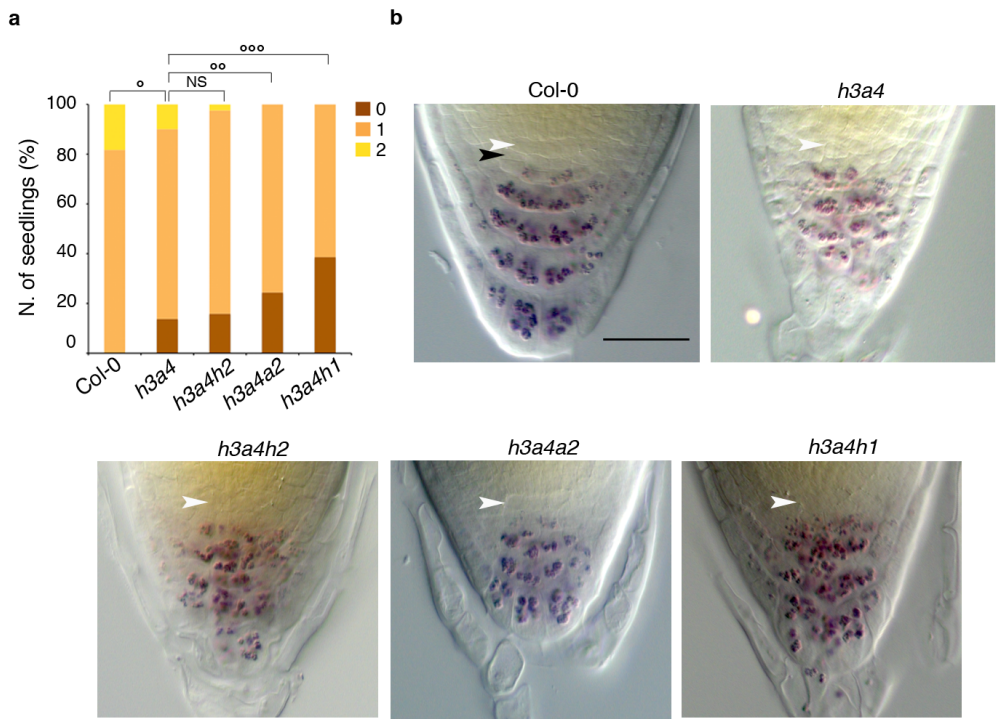

**Extended Data Fig.1. The introgression of $\gamma$ gene mutations potentiates DSCs phenotypes of *hat3 athb4* mutant. a,b, DSC phenotypes of 5d-o high order $\delta/\gamma$ mutant combinations showing enhanced CSC differentiation in *hat3 athb4 athb2* and *hat3 athb4 hat1* (*h3a4a2* and *h3a4h1*, respectively) compared with the WT (Col-0) and *hat3 athb4* (*h3a4*) and *hat3 athb4 hat2* (*h3a4h2*). Number of CSC layers (a) and representative pictures of Lugol-stained distal tissues of the indicated genotypes (b). White arrowheads indicate QC position; black arrowheads indicate CSC layers. $^{\circ}pVal_{Col-0 \text{ vs } h3a4} = 5.81 \times 10^{-4}$, $^{\circ\circ}pVal_{h3a4 \text{ vs } h3a4a2} = 0.00320$, $^{\circ\circ\circ}pVal_{h3a4 \text{ vs } h3a4h1} = 7.11 \times 10^{-5}$, $^{NS}pVal > 0.05$. Scale bar: b, 20 μm.**

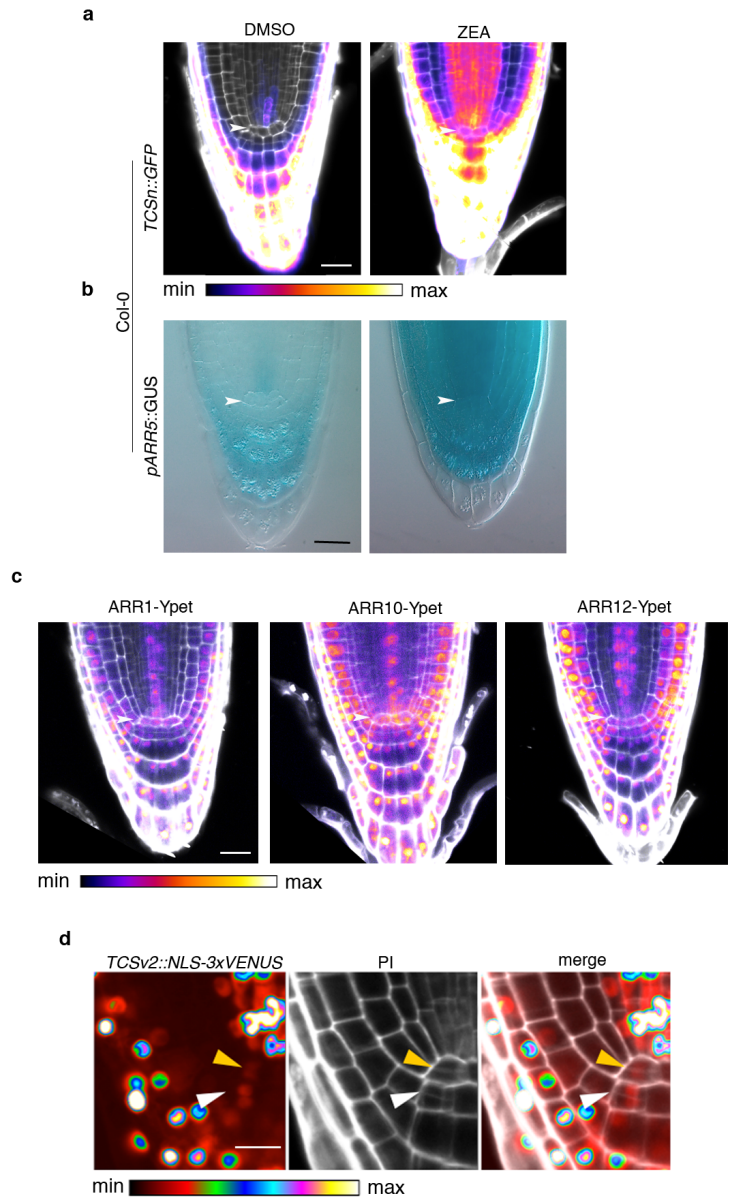

**Extended Data Fig.2. CK signaling activity in DSCs of WT seedlings. a,b,** CK signaling activation in DSCs by 24h treatments with *trans*-Zeatin (ZEA). **a,** *TCSn::GFP* expression in DSC of 5d-o seedlings untreated (DMSO) or treated with 5 μM ZEA for 24h. GFP signal intensity is displayed according to the false color scale (FCS) below, PI counterstaining is displayed in white. **b,** *pARR5::GUS* expression in DSC of 5d-o seedlings untreated (DMSO) or treated with 5 μM ZEA for 24h. Notice the lack of *TCSn::GFP* (**a**) and *pARR5::GUS* (**b**) expression in untreated DSC. White arrowheads indicate QC position. **c,** Key components of CK signaling are expressed in DSC. The images show the tissue-specific expression of *pARR1::ARR1-Ypet*, *pARR10::ARR10-Ypet*, and *pARR12::ARR12-Ypet* in 5d-o Col-0 root apices. Notice the

simultaneous presence of the three Type-B ARRs in DSCs. Ypet signal intensity is displayed according to the FCS below, PI counterstaining is displayed in white; white arrowheads indicate QC position. **d**, *TCSv2::NLS-3xVENUS* expression in dividing stem cells. The images depict the detail of a 5d-o root apex showing TCSv2 activation in dividing QC (yellow arrowhead) and CSC (white arrowhead). Left panel, *TCSv2::NLS-3xVENUS* displayed according to the FCS below; central panel, PI counterstaining; right panel, merged images. Scale bars: **a-** **c**, 20 $\mu\text{m}$; **d**, 10 $\mu\text{m}$.

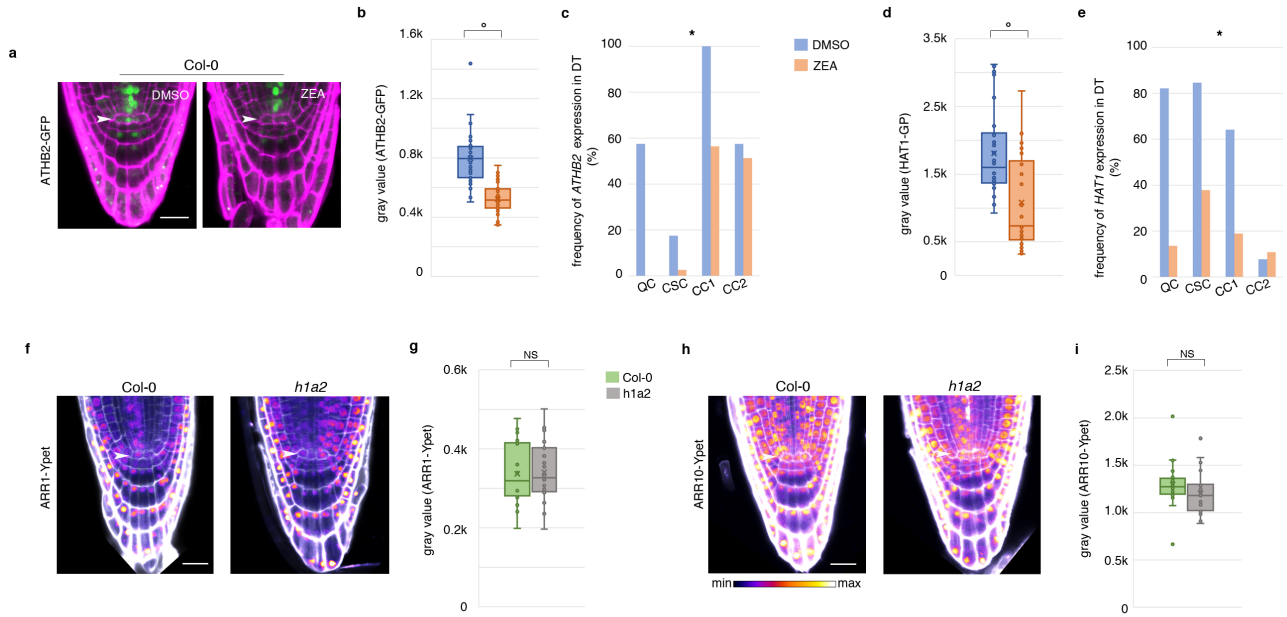

**Extended Data Fig.3. The $\gamma$ module is repressed by CK but does not affect Type-B ARRs expression.** **a-e**, ATHB2 and HAT1 are repressed by long-term CK treatments. Representative image of *pATHB2::ATHB2-GFP* seedlings grown for 5 d on control medium (DMSO) or on 0.1 μM ZEA-supplemented medium. PI counterstaining is in magenta; white arrowheads indicate QC position. **b,c**, Quantifications of ATHB2-GFP signal intensity in DSCs (**b**) and frequency of nuclear ATHB2-GFP signal in distal tissues (DT) (**c**) of roots grown as in **a**. $^{\circ}pVal_{DMSO vs ZEA} = 9.59 \times 10^{-13}$; $*pVal_{DMSO vs ZEA} = 5.50 \times 10^{-8}$. **d,e**, Quantifications of HAT1-GFP signal intensity in DSCs (**d**) and frequency of nuclear HAT1-GFP signals in DT (**e**) of roots untreated (DMSO) or treated with 0.1 μM ZEA for 5 d; refers to **Fig.3a** in the main text. $^{\circ}pVal_{DMSO vs ZEA} = 1.12 \times 10^{-4}$; $*pVal_{DMSO vs ZEA} = 3.45 \times 10^{-10}$. QC, quiescent centre; CSC, Columella Stem Cell; CC1, CSCD/Columella Cells tier 1; CC2, Columella Cells tier 2. **f-i**, the expression of Type-B ARRs is not affected in *hat1 athb2* mutants. Representative images (**f,h**) and signal intensity quantification in DSCs (**g,i**) of *pARR1::ARR1-Ypet* (**f,g**) and *pARR10::ARR10-Ypet* (**h,i**) in 5d-o Col-0 and *hat1 athb2* (*h1a2*) seedlings. Ypet signal intensity in **f,h** is displayed according to the FCS below, PI counterstaining is displayed in white; white arrowheads indicate QC position; $^{NS}pVal > 0.05$. Scale bars: **a,f,h**, 20 μm.

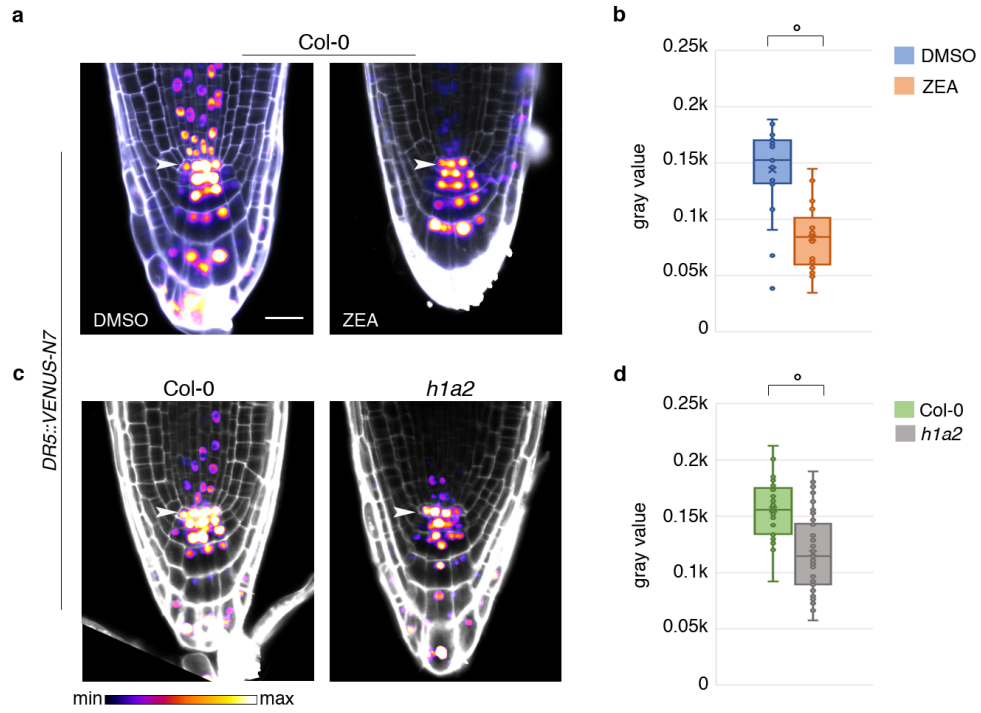

**Extended Data Fig.4. Decreased IAA signaling in CK-treated WT and *hat1 athb2* seedlings.**

**a-d**, Representative images (**a,c**) and signal intensity quantification in DSCs (**b,d**) of *DR5::VENUS-N7* in Col-0 roots untreated (DMSO) or treated with 1μM ZEA for 5d (**a,b**), and in 5d-o *hat1 athb2* (*h1a2*) seedlings compared to Col-0 (**c,d**). VENUS signal intensity in **a,c** is displayed according to the FCS below, PI counterstaining is displayed in white; white arrowheads indicate QC position. **c**, $^{\circ}pVal_{DMSO\ vs\ ZEA}=1.87\times10^{-6}$; **d**, $^{\circ}pVal_{Col-0\ vs\ h1a2}=7.65\times10^{-9}$. Scale bars: **a,c**, 20 μm.

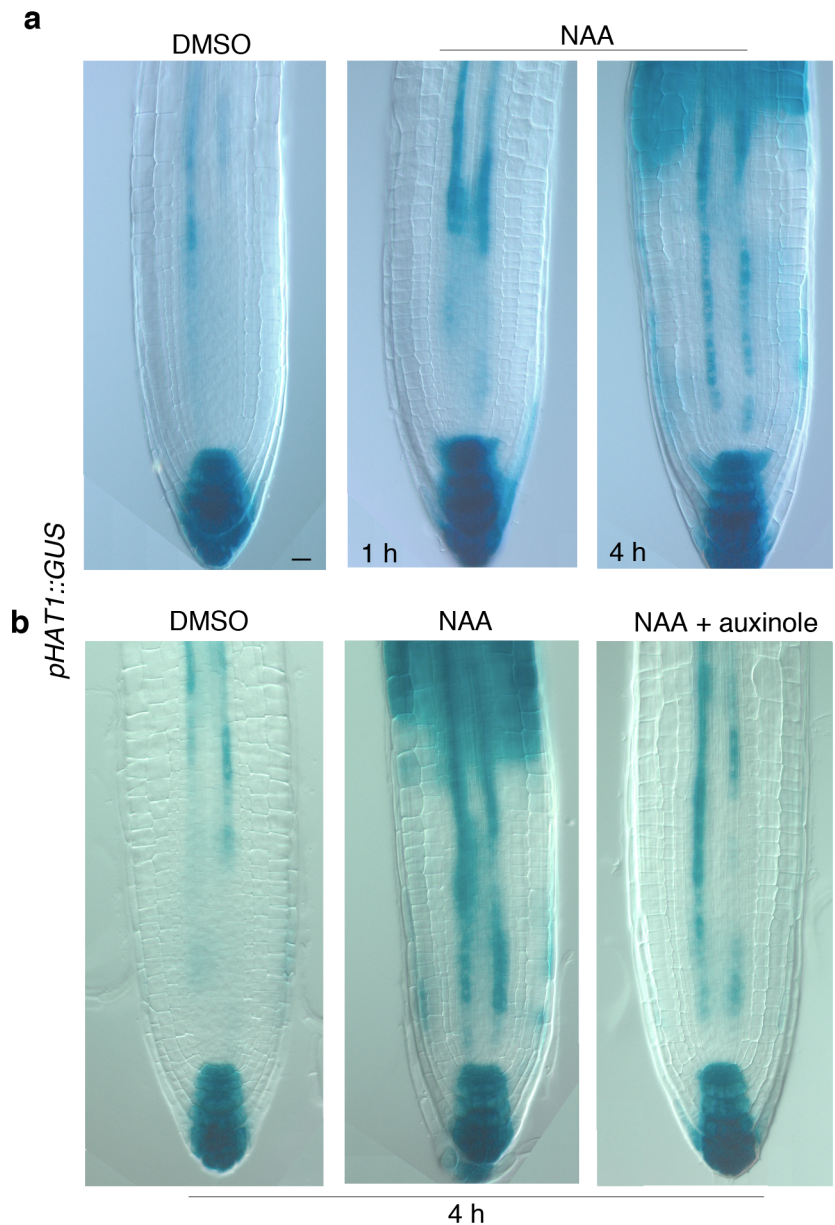

**Extended Data Fig.5. HAT1 expression is upregulated by NAA downstream of the nuclear** **auxin pathway. a,** Time-course analysis of *pHAT1::GUS* expression in 5d-o root apices untreated (DMSO) or treated with 5μM NAA for the indicated times. **b,** *pHAT1::GUS* expression in 5d-o root apices untreated (DMSO) or treated with 5μM NAA or with 5μM NAA/10μM auxinole for 4h. Notice that auxinole halts NAA-mediated induction of *pHAT1::GUS*. Scale bar: 20 μm.

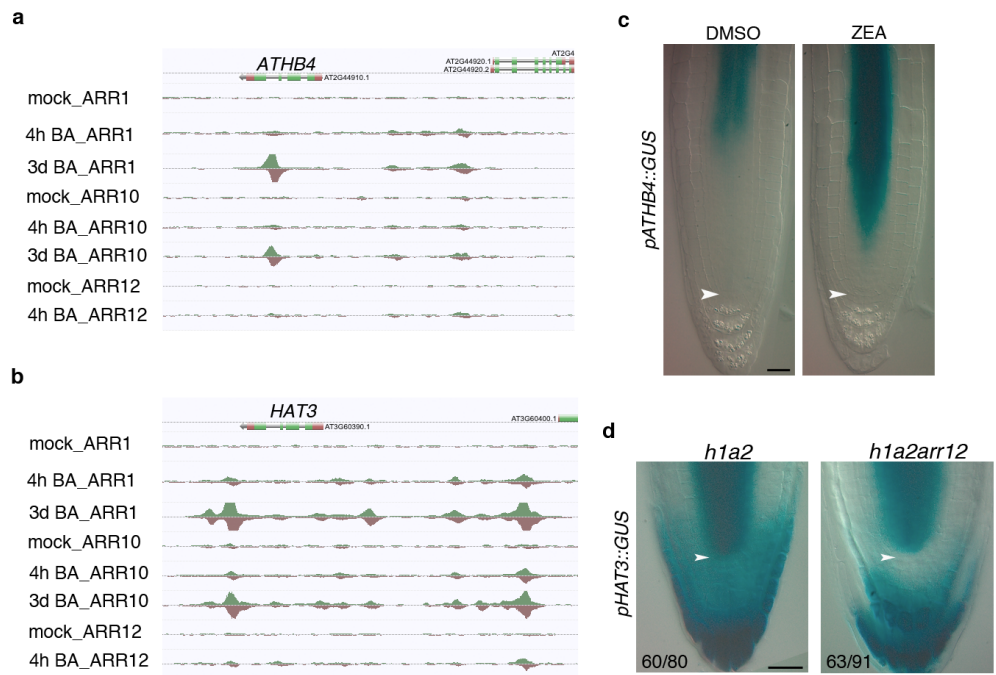

**Extended Data Fig.6. The $\delta$ module is upregulated by prolonged CK treatments. a,b,** Schematic representation of ChIP-Seq data showing enriched binding of Type-B ARR1, ARR10, and ARR12) on the genomic regions adjacent to *ATHB4* (a) and *HAT3* (b) upon prolonged CK treatment. Data were retrieved from the Salk Institute Genomic Analysis Laboratory (SiGnAL) database and refer to the dataset previously published in ref <sup>8</sup> (http://neomorph.salk.edu/aj2/pages/mxie/CK_BARR_net.php). Mock, untreated; 4h BA, 4 hours Benzyl-Adenine treatment; 3d BA, 3 days Benzyl-Adenine treatment. Notice that 3d BA data for ARR12 are not displayed because the track was not available in the database. c, *pATHB4::GUS* expression is upregulated in the vasculature of plants grown for 5d on 1μM ZEA compared to untreated control (DMSO). d, the CK-mediated induction of *HAT3* depends on ARR12. Representative images of *pHAT3::GUS* expression in DSCs of *hat1 athb2* and *hat1* *athb2 arr12* mutants. Notice the substantially reduced *pHAT3::GUS* staining in the DSCs of the triple mutant. White arrowheads indicate the QC. Scale bars: c,d, 20 μm.

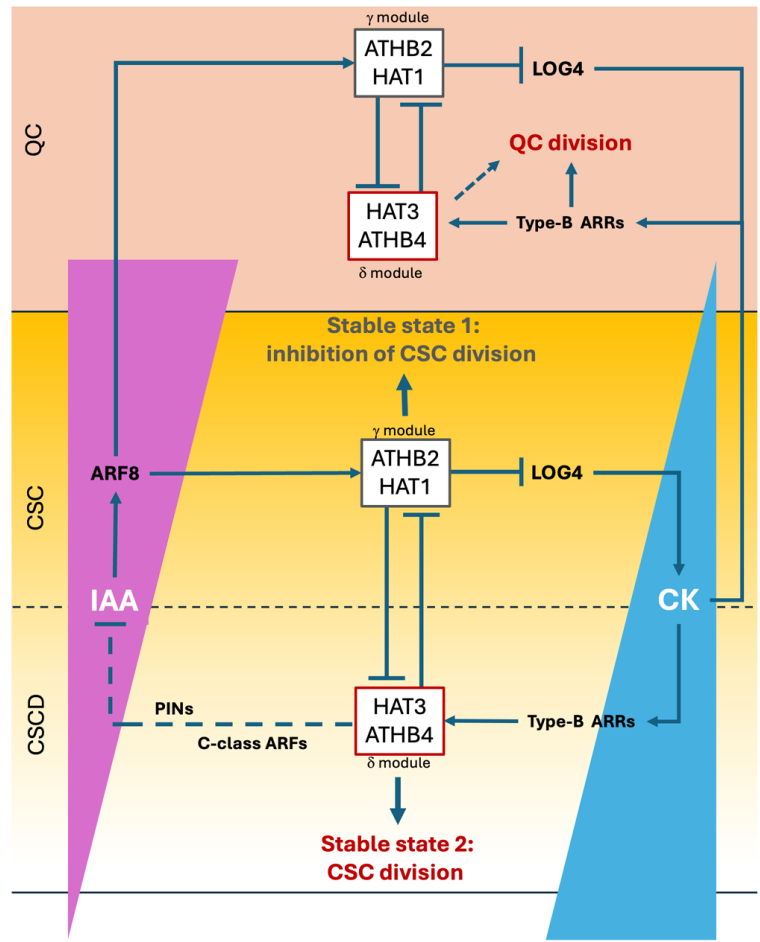

**Extended Data Fig.7. Opposing pairs of HD-Zip II TFs regulates DSC fate downstream of IAA**

**and CK behaving as a bistable switch.** Schematic model of the findings illustrated in this work.

In distal root tissues antiparallel IAA (downward) and CK (upward) signals partly overlap

across DSCs. The IAA signaling maximum sets the expression of the $\gamma$ module (HAT1 and

ATHB2) in the QC and CSCs via ARF8. In these tissues, the $\gamma$ module acts in a bipartite manner:

on the one hand, it downregulates local CK-biosynthesis via LOG4; on the other hand, it

excludes the $\delta$ module (HAT3 and ATHB4) - the actual effector of stem cell divisions – from

these tissues. By simultaneously repressing the input and the effectors of cell division, the $\gamma$

module ensures the maintenance of the DSC stemness. This configuration constitutes the

stable state 1 of the bistable switch and is represented by the presence of 1 layer of CSCs in

root apex. Upon changes of the IAA/CK ratio in DSCs, CK signaling promotes a slow

upregulation of the $\delta$ module within DSCs via Type-B ARRs. In DSCs, the $\delta$ module

downregulates the $\gamma$ module, causing a local increase of CK levels which further potentiate $\delta$

protein expression. The $\delta$ module may additionally act on IAA signaling and transport as shown in ref <sup>24</sup> (dashed line), temporarily enhancing the effect of CK. This configuration represents stable state 2 in which cell division is promoted resulting in 2 layers of CSCs. The recovery of the stable state 1 may occur through brief pulses of IAA signaling in-between CSC division events as the ones observed in ref <sup>15</sup>, which reinstate stemness and promote CSCD differentiation, maintaining the progression of root cap development. It is worth noting that within this framework CK-induced QC divisions depend only in part on the $\delta$ module (dashed arrow) and are predominantly regulated by Type-B ARRs as also described in ref <sup>30</sup>.

103 **Extended data Table 1. List of all the primers used in this study**

| **Purpose/Construct** | **Primer name** | **Sequence** |
| --- | --- | --- |
| Amplification of *HAT1* promo + CDS sequence 5' | HAT1-F | GAGCTTTTGGGTCTCGAATTTCTTCT |
| Amplification of *HAT1* promo + CDS sequence 3';<br>*XVE>HAT1* 3' | HAT1-AS-Spe I | GATCACTAGTTTAAGACCTAGGACGCATCA |
| *pHAT1::GUS* 5';<br>*pHAT1::HAT1-GFP* 5' | HAT1 -BP1 | GGGGACAAGTTTGTACAAAAAAGCAGGCTTCGATTTGGGTCTCGAATTTCT |
| *pHAT1::GUS* 3' | HAT1-BP2 | GGGGACCACTTTGTACAAGAAAGCTGGGTCCATGATATTGTCTCTTGGTA |
| *pHAT1::HAT1-GFP* 3' | HAT1-BP3 | GGGGACCACTTTGTACAAGAAAGCTGGGTCTAGACCTAGGACGCATCACAT |
| *XVE>HAT1* 5' | HAT1-S - Apa I | AATTGGGCCCATGATGATGGGTAAAGAGGA |
| Amplification of *HAT2* promo + CDS sequence 5' | HAT2-F | CGCGGTCATGCTCAATTAAGTGCTA |
| Amplification of *HAT2* promo + CDS sequence 3' | HAT2-AS | AATTACTAGTTCACGATCGTGGACGCAAGG |
| *pHAT2::HAT2-GFP* 5' | HAT2-BP1 | GGGGACAAGTTTGTACAAAAAAGCAGGCTTCGACGCGGTCATGCTCAATTAAG |
| *pHAT2::HAT2-GFP* 3' | HAT2-BP3 | GGGGACCACTTTGTACAAGAAAGCTGGGTCCGATCGTGGACGCAAGGCTT |
| **Allele genotyping** | **Primer name** | **Sequence** |
| *hat1-2* | HAT1 RP | CTCTTTAGGGGCAAAATGGTC |
|  | HAT1 LP | ACAAAGAGGGAACAAGTTGGG |
| *hat2-1* | HAT2 RP | GATCGAATGTTTGGTTCCATG |
|  | HAT2 LP | GTTGGTTGGTCCACAACTTTG |
| *hat3-3* | HAT3 RP | Described in ref.<sup>44</sup> |
|  | HAT3 LP | Described in ref.<sup>44</sup> |
| *athb4-1* | ATHB4 RP | Described in ref.<sup>44</sup> |
|  | ATHB4 LP | Described in ref.<sup>44</sup> |
| *athb2-3* | ATHB2 RP | Described in ref.<sup>44</sup> |
|  | ATHB2 LP | Described in ref.<sup>44</sup> |
| *arr1-4* | ARR1 RP | GAGATGGCATTGTCTCTGCTC |
|  | ARR1 LP | GATCAAACCCATTCAATGTCG |
| *arr12-1* | ARR12 RP | GATGACTGGCCACATTTTCTC |
|  | ARR12 LP | TTCTGGGTCTGTTGAAAAACG |
| *arf8-2* | ARF8 RP | TCGGGAGATGTCTAATGAACG |
|  | ARF8 LP | TGATGTTACAGATGGAAGGGG |
